# MSPminer: abundance-based reconstitution of microbial pan-genomes from shotgun meta-genomic data

**DOI:** 10.1101/173203

**Authors:** Florian Plaza Oñate, Emmanuelle Le Chatelier, Mathieu Almeida, Ales-sandra C. L. Cervino, Franck Gauthier, Frédéric Magoulès, S. Dusko Ehrlich, Matthieu Pichaud

## Abstract

**Motivation:** Analysis toolkits for shotgun metagenomic data achieve strain-level characterization of complex microbial communities by capturing intra-species gene content variation. Yet, these tools are hampered by the extent of reference genomes that are far from covering all microbial variability, as many species are still not sequenced or have only few strains available. Binning co-abundant genes obtained from *de novo* assembly is a powerful reference-free technique to discover and reconstitute gene repertoire of microbial species. While current methods accurately identify species core parts, they miss many accessory genes or split them into small gene groups that remain unassociated to core clusters.

**Results:** We introduce MSPminer, a computationally efficient software tool that reconstitutes Metagenomic Species Pan-genomes (MSPs) by binning co-abundant genes across metagenomic samples. MSPminer relies on a new robust measure of proportionality coupled with an empirical classifier to group and distinguish not only species core genes but accessory genes also. Applied to a large scale metagenomic dataset, MSPminer successfully delineates in a few hours the gene repertoires of 1 661 microbial species with similar specificity and higher sensitivity than existing tools. The taxonomic annotation of MSPs reveals microorganisms hitherto unknown and brings coherence in the nomenclature of the species of the human gut microbiota. The provided MSPs can be readily used for taxonomic profiling and biomarkers discovery in human gut metagenomic samples. In addition, MSPminer can be applied on gene count tables from other ecosystems to perform similar analyses.

**Availability:** The binary is freely available for non-commercial users at enterome.fr/site/downloads/ Contact

**Supplementary information:** Available in the file named *Supplementary Information.pdf*

## Introduction

Metagenomics has revolutionized microbiology by allowing culture-independent characterization of microbial communities. Its advent has allowed an unprecedented genetic characterization of the human gut microbiota and emphasized its fundamental role in health and disease (Wang et al., 2015). Shotgun metagenomics where whole-community DNA is randomly sequenced bypasses the biases and limitations of 16S rRNA sequencing (Větrovský and Baldrian, 2013; Brooks et al., 2015) by providing high resolution taxonomic profiling as well as insights into the diverse physiological roles and the metabolic potential of the community (Ranjan et al., 2016; Jovel et al., 2016).

The analysis of large cohorts revealed a substantial inter-individual microbial gene content variability (Li et al., 2014) nucleotide polymorphism (Schloissnig et al., 2012) which reflects that individuals are not only carriers of various species, but also of different strains of the same species (Greenblum et al., 2015; Zhu et al., 2015). The characterization of the accessory genes found in individual strains is crucial in many contexts as they can provide functional advantages such as complex carbohydrates metabolism (Larsbrink et al., 2014), antibiotic resistance or pathogenicity (Loman et al., 2013; Scaria et al., 2010).

Recent analysis toolkits for shotgun metagenomics data achieved strain-level resolution when coverage is sufficient. To this end, they either capture intra-species single-nucleotide polymorphisms (SNPs) in preidentified marker genes (Luo et al., 2015; Truong et al., 2017), gene content variation (Scholz et al., 2016) or both (Nayfach et al., 2016). However, these tools are hampered by the extent of the reference genomes.

Indeed, microbial variability extends far beyond the content of reference genomes making metagenomic samples an untapped reservoir of information. First, it has been estimated that on average 50% of the species present in the human gut microbiota of Western individuals lack reference genome and this proportion rises to 85% in individuals with traditional lifestyles (Nayfach et al., 2016). Even if recent advancements of culture-based methods have proven that a substantial proportion of these species are actually cultivable (Browne et al., 2016; Lagier et al., 2016), the number of unknown species is probably still important. In addition, these techniques remain laborious and time consuming. Second, although species of public health interest (e.g. *Escherichia coli, Salmonella enterica* or *Clostridium difficile)* are represented by hundreds or even thousands of strains in genome databases, only few strains are available for the great majority of commensal species. Consequently, accessory genes associated with microbial phenotypic traits may be missing in gene repertoires constructed from reference genomes.

Metagenomic assembly where overlapping reads are merged into longer sequences called contigs is a powerful reference-free technique for overcoming the limitations of reference-based methods. However, assembly remains a computationally challenging task and despite the many dedicated tools proposed, the process only recovers incomplete genomes scattered in multiple contigs (Sczyrba et al., 2017). In an attempt to obtain exhaustive references, metagenomic assembly is performed on multiple samples to create non-redundant gene catalogs (Almeida and Pop, 2015).

Then, these catalogues are used in metagenome-wide association studies for disease-related analyses (Wang and Jia, 2016) or descriptive purposes (Li et al., 2014). However, testing millions of genes is biased towards organisms with the most genes in the pool as they have more chances of being picked up. In addition, this approach lacks statistical power because many genes have strongly correlated abundances profiles which amounts to perform the same test multiple times (Schwartzman and Lin, 2011).

Considering that the physically linked genes should have proportional abundances across samples, binning co-abundant genes has been proposed to organize catalogs into clusters of genes originating from the same biological entity. However, clustering millions of genes is a computationally intensive task as pairwise comparison of all gene abundance profiles is hardly feasible. To reduce the number of comparisons, some authors have performed binning on the subset of genes that were statistically significant by themselves (Qin et al., 2012; Le Chatelier et al., 2013), which does not improve the statistical power of the analysis. Others have proposed methods to perform the clustering of complete gene references based either on the Markov clustering algorithm (Karlsson et al., 2014), the Chameleon clustering algorithm (Jie et al., 2017) or a variant of the Canopy clustering algorithm (Nielsen et al., 2014).

Although direct proportionality is expected between co-abundant genes, these methods rely either on Pearson’s or Spearman’s correlation coefficients which respectively assess a linear association with a potentially non-null intercept or any monotonic association. Thus, these coefficients are not specific enough and spurious associations can be discovered. In addition, they are hampered by rare genes with many null counts (Huson, 2007), non-normal gene counts distributions (Kowalski, 1972) and presence of outliers (Osborne and Overbay, 2004).

Furthermore, current clustering strategies group species core genes and highly prevalent accessory genes into the same cluster, but miss lower prevalence accessory genes or assign them to small separate clusters (Almeida et al., 2016). Dependency between core and accessory clusters can be evaluated downstream using the Fisher’s exact test (Nielsen et al., 2014), which compares their presence/absence patterns across samples. Yet, this strategy does not account for the co-abundance of genes and is poorly discriminative when considering accessory clusters that are rare or associated with very prevalent species. In addition, it is not suitable for detecting clusters shared between several species.

To overcome these limitations, we have developed MSPminer, the first tool that discovers, delineates and structures Metagenomic Species Pangenomes (MSPs) from large-scale shotgun metagenomics datasets without referring to genomes from isolated strains. MSPminer presents several significant improvements over existing methods. First, it relies on a robust measure of proportionality for the detection of co-abundant but not necessarily co-occurring genes as expected for non-core genes. Second, genes grouped in a MSP are empirically classified as core, accessory and shared genes.

To illustrate its usefulness, we applied MSPminer to the largest publicly available gene abundance table which is composed of 9.9M genes quantified in 1 267 human stool samples (Li et al., 2014). We show that MSPminer successfully groups genes from the same species and identifies additional genes. Gene variability of microbial species is better captured and their quantification is subsequently more precise. MSPminer is a computationally efficient multithreaded program implemented in C++ that can process large datasets with millions of genes and thousands of samples in just a few hours on a single node server.

## 1 Methods

### 1.1 Comparison of gene abundance profiles

Microbial pan-genomes are gene repertoires composed of core genes present in all strains and accessory genes present in only some of them (Medini et al., 2005). In a shotgun metagenomic sequencing context, we assumed that core genes of a microbial species should yield directly proportional mapped reads counts across samples (co-abundance) and should be consistently observed in samples if sequencing depth allows (co-occurrence). Remarkably, core genes and accessory genes should have directly proportional counts only in the subset of samples where they are both detected (**Fig. 1**). To group the core genes of a species and then identify its accessory genes, we developed a measure that evaluates proportionality between gene counts using samples where both genes are detected at a sufficiently high abundance.

**Fig. 1:**
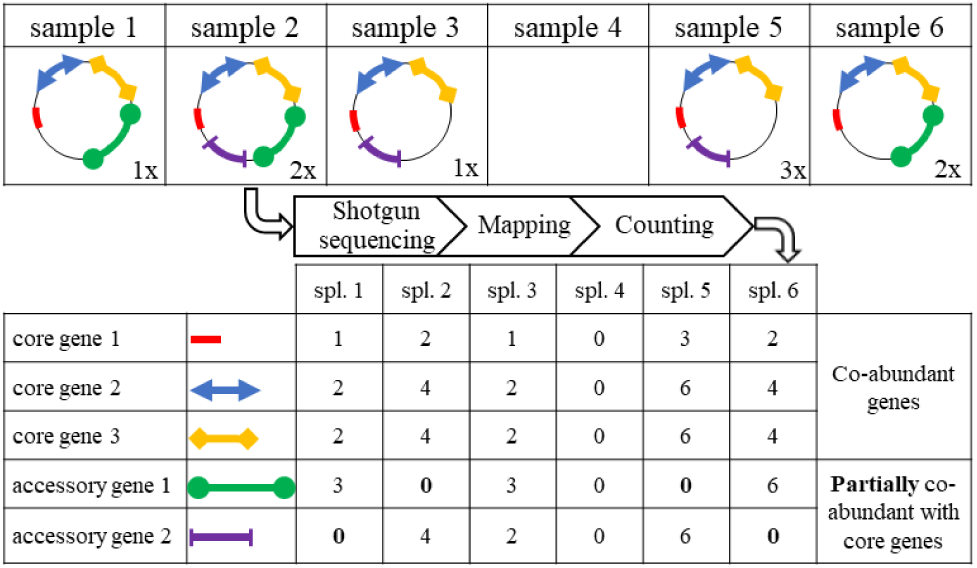
Simplified model illustrating the rationale behind the method. We consider 6 samples of which all except the fourth carry a strain of the same microbial species represented here by a circular chromosome. Each strain is composed of different genes materialized by colored circular arcs. The species genes repertoire is made up of 3 core genes and 2 accessory genes present in only some strains. Gene length based on an arbitrary scale equals 1 (core gene 1), 2 (core gene 2, core gene 3 and acc. gene 2) or 3 (acc. gene 1). A shotgun sequencing experiment is performed on each sample with a read length of 1 (length of the shortest gene). The sequencing coverage is indicated at the bottom right of the chromosome of each strain. Finally, a table counting the number of reads aligned on each gene in each sample is generated. In a given sample, the number of reads aligned on a gene is equal to its length multiplied by the sequencing coverage. A directly proportional relationship is observed between the abundance profiles of core genes, the proportionality coefficient being equal to the ratio of their length. In contrast, such relationship between a core and an accessory gene is observed only in the subset of samples where the accessory gene is present.

#### Estimation of the coefficient of proportionality

Let *S* = {*s*_1_, *s*_2_, …, *s_m_*} be a set of *m* metagenomic samples. Let 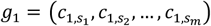 and 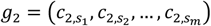 be the vectors of the number of mapped reads on the two genes to be compared.

Suppose there is a relationship of proportionality between *g*_1_ and *g*_2_ noted *g*_2_ = *α* · *g*_1_. Here, the coefficient of proportionality *α* is a strictly positive constant roughly equal to the ratio of *g*_2_ and *g*_1_ length. However, this ratio is not a good estimator when genes are duplicated or when their coverage is not uniform (**Supplementary Fig. 1**). Therefore, *α* was robustly estimated with the following iterative algorithm:

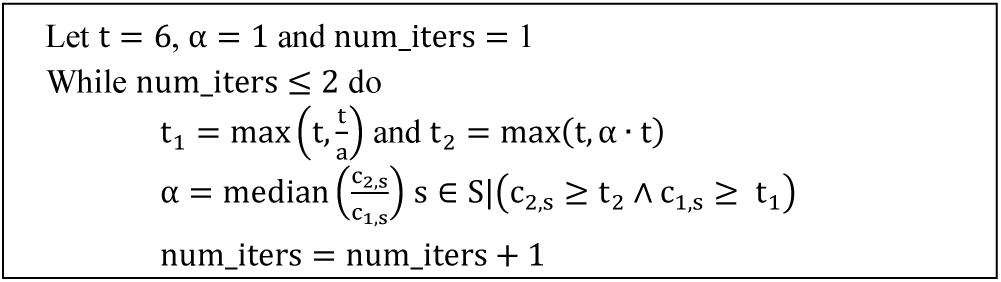

When estimating *α*, only samples were *g*_1_ and *g*_2_ had counts above a threshold *t* were taken into account. In the second iteration, different quantification thresholds named t_1_ and t_2_ were used to reflect that one gene may have higher counts than the other. This filtering has the following advantages:

1. It discards samples where both genes have null counts as they do not provide a quantitative information which can be used for the assessment of proportionality.
2. It discards samples with low counts which do not allow a precise estimation of the coefficient of proportionality.
3. It discards samples where one of the genes has a null count to detect proportionality occurring in a subset of samples only.

If less than 3 samples were available for the estimation of α, the genes were not compared.

#### Classification of zeros

Quantification thresholds were also used to classify zeros. A gene with a null count in a sample can be either a sampling or a structural zero. In the first case, the gene is not detected because of sampling or technical effects while in the second case the gene is really absent in the sample. Distinguishing these two kinds of zeros is crucial to accurately classify a gene as core or accessory.

Here, a gene with a null count in a sample was classified as a structural zero if the other gene had a count above its quantification threshold i.e. (c_2_,_s_ ≥ t_2_ Λ c_1,s_ ≥ t_1_) or (c_2_,_s_ = 0 Λ c_1,s_ ≥ t_1_). Otherwise, it was classified as an undetermined zero.

With an initial threshold t = 6, the probability of misclassifying a null count as a structural zero is 0.2% under the assumption that the number of reads mapped on a gene follows a Poisson distribution (*P*(*X* = 0|λ = 6) = 0.002).

#### Non-robust measure of proportionality

First, counts were square root transformed to stabilize variance and reduce the skewness of their distribution (Bland and Altman, 1996). Then, the directly proportional relationship between the two genes was evaluated by a modified version of the Lin’s concordance correlation coefficient (Lin, 1989):

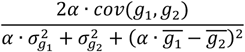
where 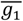 and 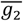 are the means, 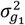 and 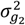 are the variances and *cov*(*g*_1_, *g*_2_) is the covariance of *g*_1_ and *g*_2_. Only samples where both genes had non-null counts were considered to compute this coefficient.

#### Robust measure of proportionality

We derived a robust version of the measure to identify associated genes despite the presence of samples with inconsistent counts, hereafter named outliers. Indeed, outliers may decrease significantly the concordance coefficient calculated if not accounted for.

Residuals defined as the difference between observed and expected proportional counts were computed on samples where both genes had counts above their respective quantification thresholds with the following formula:

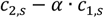

Then outliers were detected using the Tukey’s method. Let *Q*_1_ and *Q*_3_ be the first and third quartiles of the residuals and *IQR* be the interquartile range defined by *IQR* = *Q*_3_ − *Q*_1_. Among the *m*′ samples with non-null counts in both genes, those with residuals greater than *Q*_3_ + *I*. 5 · *IQR* or lower than *Q*_1_ − *I*. 5 · *IQR* were classified as outliers.

Finally, the measure of proportionality was computed on remaining samples. To avoid the detection of spurious associations with too many outliers, this robust measure was not computed if there were more than (*m*′ − 5) · 0.3 outliers that is to say a percentage of asymptotically equal to 30%.

### 1.2 Reconstitution of Metagenomic Species Pan-genomes

We developed MSPminer, a method that uses the measures of proportionality described above to group co-abundant genes into <u>M</u>etagenomic <u>S</u>pecies <u>P</u>an-genomes (MSPs). MSPminer empirically distinguishes core from accessory genes based on their presence absence patterns and tags genes observed in samples where the core is not detected as shared.

MSPminer is implemented in C++ and uses the OpenMP framework to take advantage of multicore processors.

#### Input data and filtering

MSPminer takes as input a tab-separated values matrix giving the number of reads mapped on *n* genes (rows) across *m* metagenomic samples (columns). Count data is neither normalized by gene length, nor by read length nor by sequencing depth. Indeed, the number of times a gene is detected is lost after normalization while it is used to classify null counts.

Rare genes which do not support enough quantitative information for further processing were discarded. By default, genes with counts greater than 6 in at least 3 samples were kept.

#### Gene binning

To avoid comparison of all pairs of genes, genes with the highest count in the same sample were binned. On real metagenomic data, we found that this strategy not only decreases the number of comparisons to perform but increases the probability that related genes are placed in the same bin compared to random assignment (**Supplementary Fig. 2**).

To achieve a good load balancing, raw read counts were normalized prior to bin assignment by the number of mapped reads in samples, as samples with high sequencing depth tend to bin more genes. (**Supplementary Fig. 3**). Normalized counts were used in this step only.

#### Seeds creation

Sets of co-abundant and co-occurring genes called *seeds* hereafter were identified. Seeds were created in parallel in each bin by a greedy approach. First, genes were compared pairwise. All pairs of genes with a non-robust measure of proportionality of at least 0.8 and no structural zeros were saved in a list. Then, the list was sorted by decreasing measure of proportionality.

The pair of genes with the highest measure of proportionality was selected as a centroid. Genes related to one of the centroid genes were grouped together in a new seed and removed from the list. This procedure was repeated until the list was empty.

#### Seed representative

For each seed, a pseudo gene referred as *representative* was computed to compare seeds with each other. First, the seed representative was defined as the median vector of the counts of all its genes. Then, each gene of the seed was compared to the seed representative using the measure proportionality. The final seed representative corresponded to the median vector of the counts of the 30 genes with the highest measure of proportionality.

#### Seeds merging across bins

Some related genes may have been assigned to different bins when samples with the highest counts had close values. Therefore, seeds with a non-robust measure of proportionality of at least 0.8 and no structural zeros counts were merged. After merging, seeds with less than 150 genes were discarded.

#### Core seeds identification

Core seeds were identified among final seeds, based on the assumption that in a set of related seeds, the largest corresponds to a species core genome and the others are modules of either accessory or shared genes.

To this end, seeds were sorted by decreasing number of genes. The largest seed was defined as a new core seed. Then, the representative of the core seed was compared to the representative of all remaining seeds. Seeds with a robust measure of proportionality of at least 0.8 with the core seed were discarded from the list of potential cores. The procedure was iterated until there was no more seed to process

#### Identification of genes associated to core seeds

The representatives of each core seed were compared to all the genes. Genes with a robust measure of proportionality of at least 0.8 were considered as associated to the core seed.

#### Classification of genes associated to a core seed

Let *g*_1_ be the median vector of the number of mapped reads on a core seed and *g*_2_ the vector of the number of mapped reads on a gene related to this core seed. The related gene was assigned to one of the 4 following classes according to the presence of structural zeros:

1. core: the related gene was present in all the samples where core seed was detected and uniquely in those (**Fig. 2.A**)

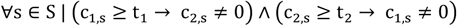
2. accessory: the related gene was present in a subset of samples where core seed was detected (**Fig. 2.B**)

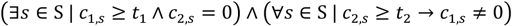
3. shared core: the related gene was detected in all the samples where the core seed was present plus some samples where the core seed absent (**Fig. 2**.**C**)

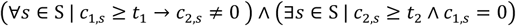
4. shared accessory: the related gene was detected in a subset of samples where the core seed was present plus some samples where the core seed was absent (**Fig. 2.D**)

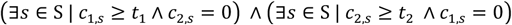

**Fig. 2:**
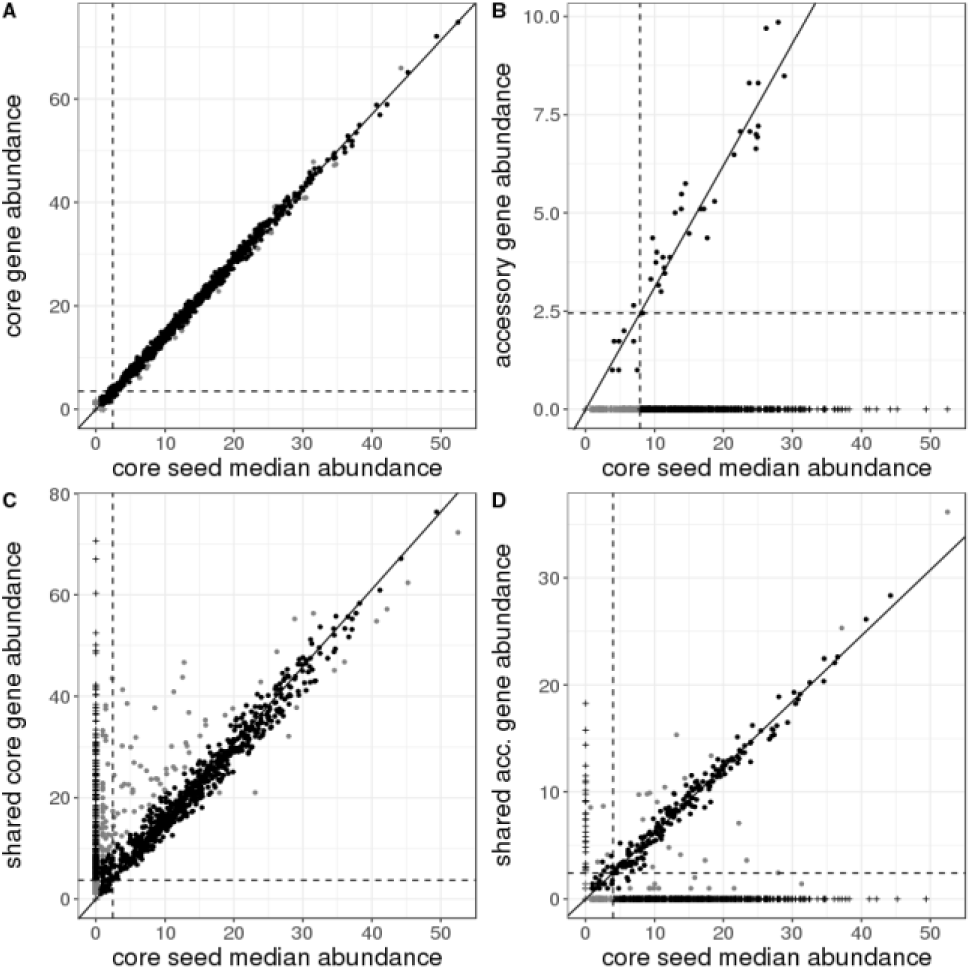
Illustration of the four types of genes grouped in a MSP. The median abundance profile of the 30 best representative genes of a core seed (x-axis) is compared to the abundance profile of four of its related genes (y-axis). Genes are quantified across the 1267 samples of the IGC catalog. Abundances are represented on a square root scale. The slope of the solid line is equal to *α*. The intercepts of the vertical and horizontal dashed lines are respectively *t*_1_ and *t*_2_. Black and grey points are respectively inlier and outlier samples. Black and grey crosses on the x and y axes are respectively structural and undetermined zeros. Only structural zeros are taken into account to affect genes to a given class. **A**. The gene is classified as core. It is present in all the samples where core seed is detected and only in those. **B**. The gene is classified as accessory. It is present in a subset of samples where core seed is detected (7.2%). **C**. The gene is classified as shared core. It is present in all the samples where core seed is detected plus 286 samples where the core seed is absent **D**. The gene is classified as shared accessory. It is present in a subset of samples where core seed is detected (30.6%) plus 28 samples where the core seed is absent.

#### Creation of Metagenomic Species Pan-genomes

Core, accessory, shared core and shared accessory genes associated to a core seed were assembled in a MSP.

Core genes were compared to the core seed representative and sorted by decreasing measure of proportionality. In each class except core, a clustering procedure similar to the one used to create seeds was run. It identified modules of co-occurring genes that may be interpreted as functional units, i.e. operons. Unclustered genes were saved as singleton modules.

### 1.3 Simulated dataset

For evaluation purposes, we generated abundance tables simulating the counts of genes from a single virtual species. The pan-genome of this species consisted in 1 000 core genes detected in all strains and 6 000 accessory genes present only in some of them. Gene lengths were randomly drawn between 100 bp and 5 000 bp. The prevalence of accessory genes was randomly drawn between 2.5% and 99.5%.

In a first simulation, 200 samples containing each a single strain of the species were generated. The sequencing coverage of a strain in a sample was drawn from a uniform law (min=0.6, max=20) and read length was set to 100 bp. In a given sample, the theoretical number of reads mapped on a gene was calculated according to the gene length, the strain coverage and the presence or not of the gene in the strain. Finally, the observed gene counts were drawn from Poisson distributions with means equal to theoretical counts.

In the second simulation used to evaluate the robust measure, outliers were added by multiplying observed counts of each gene by either ¼, ⅓, 2, 3 or 4 in 5%, 10% and 20% of the samples were it was present.

Next, we progressively decreased the number of samples where the species was detected (200, 100 and 50) to apprehend the impact of this parameter on the completeness of MSPs.

Finally, we simulated samples carrying two strains of the species where the dominant strain is 5 to 10 times more abundant than the subdominant one as observed in fecal samples (Truong et al., 2017).

## 2 Results

### 2.1 Evaluation on simulated data

#### Evaluation of the measures of proportionality

First, we simulated the abundance table of a species across 200 samples to compare the performance of Pearson’s correlation coefficient, Spearman’s correlation coefficient and the proposed measure of proportionality for detecting a relation between the abundance profile of the species core genome and all its genes including accessories. Pearson’s and Spearman’s correlation coefficients decreased with the prevalence of the tested gene, while the proposed measure remained high, as it only uses samples where both the species core and the tested gene are detected (**Fig. 3.A**). Therefore, the association between core genes and many accessory genes will be missed using the correlation coefficients. However, accessory genes observed in similar subsets of samples could be grouped into small distinct clusters as their abundance profiles should be correlated. Our simulations also show that the measure of proportionality is more sensitive to species with highly variable coverage and on long genes as their counts are higher and less dispersed (**Supplementary Fig. 4**).

Then, we compared the robust measure of proportionality against its nonrobust counterpart by adding an increasing percentage of outliers to the genes abundance profiles. For a given percentage of outliers, each of these genes was compared to the outlier-free abundance profile of the core. This simulation showed that the non-robust measure of proportionality decreases when the percentage of outliers increases whereas the robust measure remains high, demonstrating that proportionality is still detected (**Fig. 3.A**).

**Fig. 3:**
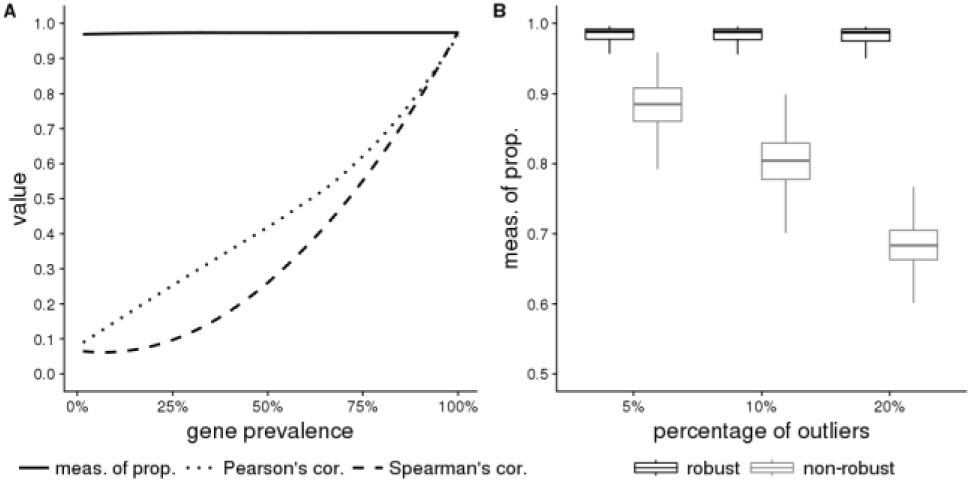
Evaluation of the measures of proportionality. **A**. Comparison of the Pearson’s correlation coefficient, the Spearman’s correlation coefficient and the proposed measure of proportionality to detect an association between the median abundance vector of the core genes of the simulated species and the abundance vectors of each of its genes. The x-axis corresponds percentage of samples where a gene is detected and the y-axis corresponds to the intensity of the relationship between the compared vectors. The closer the value is to 1, the stronger the intensity of the relationship. **B**. Comparison of the performances of the robust (black) and the nonrobust (grey) measures of proportionality to detect a relationship between the noisy abundance vector of each gene of the simulated species and the outlier-free median abundance vector of its core genes. The proportion of outliers is gradually increased to 5%, 10% and 20%.

#### Evaluation of the clustering algorithm

Next, we tested if the number of samples where the species was detected had an influence on the completion of its corresponding MSP. Although this parameter did not impact the clustering of core and prevalent accessory genes, rarer accessory genes were grouped in the MSP only when the species was detected in a sufficiently large number of samples (**Fig. 4**).

**Fig. 4:**
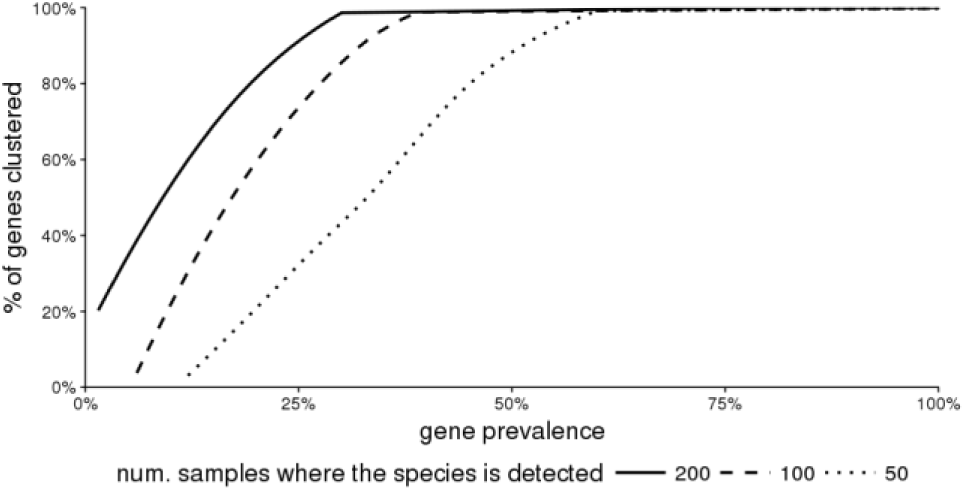
Impact of number of samples where the simulated species is detected on clustering. Finally, we explored the impact of mixture of multiple strains of the same species in samples. When occasional, strains mixture had little impact on clustering. If it was more frequent, many accessory genes of low or medium prevalence were missed (**Fig. 5**). However, strains mixture might have less impact on the clustering performance. When it occurred, we considered that the presence of a gene in one strain was independent of its presence in the other. Yet, the low nucleotide divergence frequently observed between strains present in the same fecal sample suggests that they may have similar gene content (Truong et al., 2017).

**Fig. 5:**
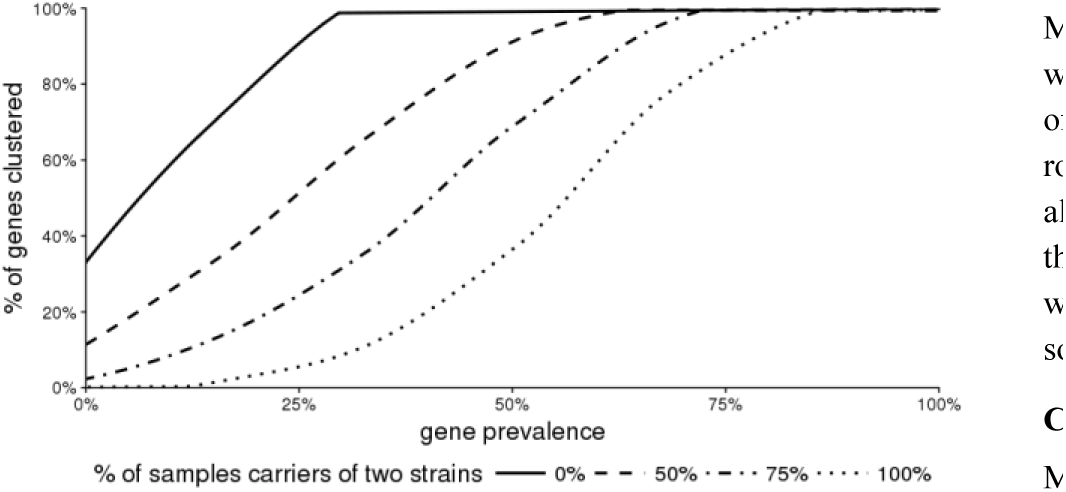
Impact of strain mixture on clustering

### 2.2 Application to the study of the human gut microbiota

We applied MSPminer to the largest publicly available gene abundance table provided with the Integrated Gene Catalog of the human gut microbiome (Li et al., 2014). In this table, 9 879 896 genes are quantified across 1 267 stool samples from individuals of various geographical origin (Europe, USA and China) and diverse health status (healthy, obese, diabetic, with inflammatory bowel disease etc.). 6 971 229 (70.6%) genes with counts greater than 6 in at least 3 samples were kept. Among these, 3 288 928 (47.2%) were organized into 1 661 MSPs (**Supplementary Table 1**).

#### Census of universal single copy marker genes

To check that MSPs correspond to real microbial species and evaluate the completeness of their core genomes, we identified 40 universal single copy marker genes (SCM) in the gene catalog (Sunagawa et al., 2013). 84% of SCMs detected in at least 3 samples were assigned to MSPs, indicating that MSPs capture a large proportion of the biological signal at species level. 915 MSPs (55%) had at least 30 SCM and 406 (24%) had all of them (Supplementary Table 2). As housekeeping genes, SCMs are essential to the microbe survival and expected among core genes. Indeed, 93% of the SCMs were core genes in their respective MSP and 70% of non-core SCMs were accessory genes of high prevalence (≥ 90%). This shows that the heuristic used for the classification of genes is reliable.

#### Precision

We evaluated the precision of MSPminer by calculating in MSPs the fraction of genes assigned to the dominant species (**Supplementary Table 4.A**). Apart from unassigned genes, the taxonomic consistency was very high for all gene categories (mean > 98%) except shared accessory genes (mean = 83.3%). Remarkably, some MSPs such as those representative of *Bacteroides plebeius*, *Ruminococcus bicirculans* and *Eubacterium eligens* had many unknown accessory genes (resp. 2 888, 2 821 and 2 399) which is coherent with the low number of genomes available for these species. On average, 80% of these novel accessory genes were validated by performing the taxonomic annotation of the contigs they derived from. The remaining genes were found in unassigned contigs or contigs carrying only one gene. Conversely, 99% of the genes of the MSP representative of *Escherichia coli* (msp_0005) were annotated as thousands of references are available for this species.

#### Sensitivity

Then, we aligned 3 143 genomes representative of 322 species of the human gut microbiome against the IGC catalog. For each genome, we defined the sensitivity as the number of its genes grouped in the most representative MSP divided by the total number of its genes found in the catalog (**Supplementary Table 4.C**). Overall, the sensitivity weighted by the number of genomes per species was high (median=77%). Interestingly, genes grouped in MSPs were significantly longer than those that were not (median length of 975 bp vs 670 bp, Wilcoxon rank-sum test p-value = 0).

More specifically, genomes of 1 127 human gut-associated *E. coli* strains were well covered by the msp_0005 (mean = 83.4%). 95% of core genes of genomes were also tagged as core in the MSP which shows again the robustness of the classification. However, 32 078 genes from the IGC catalog detected in *E. coli* genomes were missing in the msp_0005. 85% of these genes were present in less than 5% of the metagenomic samples where *E. coli* was detected, indicating that MSPminer misses rarest accessory genes which can be very numerous.

#### Comparison to the Canopy clustering algorithm

MSPminer was compared to the Canopy clustering algorithm (Nielsen et al., 2014) which is the only gene binning tool publicly available. Both tools were applied to the metagenomic dataset described above using default parameters (**Supplementary Methods**). In total, MSPminer grouped 17.8% more genes than Canopy (3 288 928 vs 2 704 552) although MSPminer had a more stringent gene selection criterion (6 971 229 vs 7 304 439 genes processed). Both tools had a very high precision (mean > 98%) but MSPminer brought a significant gain in sensitivity (median: 77% vs 62%) (**Supplementary Table 4**). Remarkably, Canopy produced more objects with at least 150 genes than MSPminer (2 010 CAGs vs 1 661 MSPs) as it splits some species (e.g. *E. coli*) into multiple clusters. In contrast, MSPminer generated one MSP per species which improves downstream statistical analysis. Finally, MSPminer achieved better computing performance than Canopy (wall time: 2h 40min vs 42h) while consuming less memory (peak memory: 74Go vs 231Go).

#### Taxonomy

642 MSPs (38.7%) could be annotated at species level, 315 (19.0%) at genus level, 525 (31.6%) at a higher taxonomic level from family to superkingdom and the remaining 179 (10.8%) could not be annotated, indicating that a majority of MSPs correspond to species not represented in GenBank (**Supplementary Fig. 5** and **Supplementary Table 3.C**). Among the annotated MSPs, one corresponded to *Homo sapiens*, 4 were unicellular eukaryotes of the genus Blastocystis, 8 were Archaea and the remaining 99% were Bacteria represented predominantly by the phyla Firmicutes (1 016 MSPs), Bacteroidetes (263 MSPs), Proteobacteria (94 MSPs) and Actinobacteria (46 MSPs).

Among the 642 MSPs annotated at species level, 304 corresponded to well-defined species and 338 matched genomes with imprecise taxonomy (i.e. *sp., cf., CAG* or *bacterium*). In the end, most MSPs assigned to well-defined species matched RefSeq reference genomes.

Interestingly, 15 species were represented by multiple MSPs such as *Faecalibacterium prausnitzii* (7 MSPs), *Bacteroides fragilis* (2 MSPs) or *Methanobrevibacter smithii* (2 MSPs) (**Supplementary Table 3.D**). In these cases, one of the MSPs matched the species reference genome and the other MSPs matched other genomes only. The low Average Nucleotide Identity (ANI) between these genomes and the species reference suggests that they actually belong to distinct species.

Conversely, 8 MSPs were attributed to reference genomes of different species (**Supplementary Table 3.E**). For all cases, the comparison of the reference genomes revealed an ANI > 96%, suggesting that they actually belonged to the same species despite distinct names were attributed.

Among the 3 813 genomes that matched MSPs annotated at species level, 369 with imprecise taxonomy could be reassigned to well-defined species, and 581 appeared misannotated or contaminated (**Supplementary Table 3.B**).

#### MSPs content

Most MSPs were small (median number of genes = 1 821) even if 51 had more than 5 000 genes (**Supplementary Fig. 6** and **Supplementary Table 2**). As expected, a strong positive correlation (Pearson’s r = 0.78) between the total number of genes in a MSP and its number of accessory genes was observed. Interestingly, 4 outliers corresponding to the unicellular eukaryotes previously described had a high number of core genes and few accessory genes. This suggests that Eukaryotic genomes have a larger number of genes and a lower gene content variability than Prokaryotes. Among the MSPs with the more accessory genes, many corresponded to species reported as highly variable such as *Klebsiella pneumoniae* (Holt et al., 2015) or *Clostridium bolteae* (Dehoux et al., 2016). As previously observed in population genomics studies comparing multiple strains of the same species (Koonin and Wolf, 2008), the prevalence of accessory genes in MSPs often follows a bimodal distribution showing either a high or low prevalence but rarely intermediate (**Supplementary Fig. 7**).

#### MSPs prevalence

Most MSPs were very rare as 596 (35.9%) were detected in less than 1% of samples and 1 110 (66.2%) in less than 5%. Only 82 (4.9%) MSPs were detected in at least half of the samples showing that the common microbial core of the human gut microbiota is limited to a few dozen species (**Supplementary Table 2**). MSPs annotated at species level were significantly more frequent than those with less precise annotation (median prevalence: 5.4% vs 1.7%, p-value=1.4.10^−21^ Wilcoxon rank-sum test) indicating that non-sequenced species are generally rarer. No clear relation between the prevalence of the MSPs and their mean abundance was found. However, 2 MSPs corresponding to *Bacteroides vulgatus* and *Bacteroides uniformis* were both very prevalent (detected in 97.5% and 94.0% of the samples respectively) and very abundant (mean relative abundance of 7.3% and 4.1% respectively). Interestingly, many rare MSPs assigned to the *Prevotella* genus were abundant in the few samples which carried them.

#### MSPs quantification for biomarkers discovery

To demonstrate that MSPminer was useful for biomarkers discovery, we first looked for differentially abundant MSPs according to the geographical origin of samples (**Supplementary Methods**). We discovered 343 MSPs differentially abundant between Westerners and Chinese including 259 more abundant in Westerners and 84 in Chinese (**Supplementary Table 5.A**). Among the discriminant MSPs, all those assigned to the Proteobacteria phylum (*Klebsiella pneumoniae*, *Escherichia coli* and *Bilophila wadsworthia*) were more abundant in Chinese which is consistent with previously published results (Li et al., 2014). Interestingly, three MSPs assigned to *Faecalibacterium prausnitzii* were significant but two were more abundant in Westerners and the other in Chinese. In addition, we discovered 134 MSPs differentially abundant between Europeans and Americans of which 119 were more abundant among Europeans (**Supplementary Table 5.B**). This result is consistent with previous studies showing lower gut microbiota diversity among Americans compared to Europeans (Sunagawa et al., 2013).

Secondly, we used MSPs for strain-level analysis. To do this, we looked for accessory genes more frequent in samples of a given geographical origin (**Supplementary Methods**). We found 51 MSPs with at least 200 such accessory genes (**Supplementary Table 5.C**). Some MSPs contained genes associated with sample origin while the abundance of their core was not, illustrating the complementarity of the two approaches.

## 3 Discussion

### 3.1 Identification of genes with proportional counts

MSPminer relies on a new robust measure to detect genes with directly proportional counts. This relation more stringent than those assessed by Pearson’s or Spearman’s correlation coefficients was successfully used to reconstitute Metagenomic Species Pan-genomes of the human gut microbiota. In fact, most genes from sequenced genomes were grouped into a single MSP showing that direct proportionality is the most common relation between genes from the same biological entity.

However, MSPminer misses some genes for which counts are not ruled by this relation. Indeed, proportionality is disrupted when gene copy number varies across samples (Greenblum et al., 2015), when a sample contains multiple strains of the same species (Truong et al., 2017), when a gene is subject to horizontal gene transfer (Dagan et al., 2008) or when genes from different species have the same reference in the gene catalogue. Nevertheless, the first two cases have most likely a limited impact as the majority of strains tend to have the same gene copy numbers (Greenblum et al., 2015) and samples often carry a dominant strain (Truong et al., 2017). Regarding shared genes, their signals are a linear combination of the MSPs that carry them. Thus, they will be identified only if these MSPs are mostly detected in separate sets of samples.

### 3.2 Parameters impacting the quality of the MSPs

The quality of the MSPs is impacted by the upstream steps required for generating the count matrix, as well as by the biological and ecological characteristics of the dataset. At the sequencing level, the number of reads (sequencing depth) generated for each sample impacts the detection and coverage of subdominant species, while read length affects the quality of the assembly and the ability to assign a read to a gene without ambiguity. At the bioinformatics level, assembly, gene prediction, gene redundancy removal, mapping and counting require expertise to select the most appropriate strategies, tools and parameters. Indeed, assemblers returning chimeric contigs which combine sequences from highly related species, inaccurate predictors generating truncated or merged genes, redundancy removal with a common threshold for all genes (95% of nucleotide identity) lead to genes of variable quality in catalogues. When quantifying genes, keeping only uniquely mapped reads underestimates the abundance of some genes whereas considering shared reads can generate false positives. As shown on simulated data and verified on a real metagenomic dataset, longer genes are more likely to be clustered in MSPs because they have greater and less dispersed counts. Finally, at the biology level, a high number of samples with varied phenotypes will improve the comprehensiveness and quality of MSPs. Indeed, as the number of samples grows, MSPminer will be able to identify rare species and assign rarer accessory genes to their respective MSPs. In addition, highly prevalent accessory genes will be reclassified from core to accessory as observed while sequencing an increasing number of strains of a species (Touchon et al., 2009).

### 3.3 Applications

As illustrated in this paper, MSPs can be used for taxonomic profiling of human gut metagenomes. By using a dedicated pipeline (Kultima et al., 2012; Karlsson et al., 2014), the sequencing reads need to be mapped on the IGC catalog to get the number times each gene was sequenced. Then, the aggregation of the core genes abundance profiles of each MSP allows accurate detection and quantification of microorganisms in samples up to species level. New MSPs will need to be built if those provided are not representative of the studied ecosystem.

Compared to methods relying on reference genomes (Truong et al., 2017), information from unknown or non-sequenced species can be exploited. In addition, our method is not impacted by contaminated genomes or incorrect taxonomic annotation. Compared to methods quantifying a few dozen universal marker genes (Sunagawa et al., 2013), MSPminer may improve the estimation of species abundance by automatically detecting among hundreds of core genes those with the highest specificity, the highest counts and lowest dispersion.

Furthermore, in each MSP, accessory genes associated with the tested phenotype can be explored opening the way to global strain-level analyses. This allows the comparison of strains carried by individuals and discovery of biomarkers corresponding to functional traits specific to certain strains.

Finally, MSPminer provides microbial population genetics from large cohorts which can help culture-dependent methods prioritize species of greater interest, such as those with no reference genome available or with reference genomes distant from the strains present in metagenomic samples (Fodor et al., 2012). When sequencing coverage allows, genomes of these species can be directly reconstituted from metagenomic assemblies by binning contigs carrying genes of the same MSP.

## Funding

This work was funded by Enterome, the ANRT (Association Nationale de la Recherche et de la Technologie) via the grant CIFRE 2014/0057 and INRA Meta-GenoPolis via the grant “Investissements d’avenir” ANR-11-DPBS-0001.

## Conflict of Interest

none declared.

## References

Almeida, M. et al. (2016) Capturing the most wanted taxa through cross-sample correlations. ISME J., 10, 2459–2467.

Almeida, M. and Pop, M. (2015) High-Throughput Sequencing as a Tool for Exploring the Human Microbiome Elsevier Inc.

Bland, J.M. and Altman, D.G. (1996) Statistics Notes: Transforming data. Bmj, 312, 770–770.

Brooks, J.P. et al. (2015) The truth about metagenomics: quantifying and counteracting bias in 16S rRNA studies. BMC Microbiol., 15, 66.

Browne, H.P. et al. (2016) Culturing of ?unculturable? human microbiota reveals novel taxa and extensive sporulation. Nature, 533, 543–546.

Le Chatelier, E. et al. (2013) Richness of human gut microbiome correlates with metabolic markers. Nature, 500, 541–546.

Dagan, T. et al. (2008) Modular networks and cumulative impact of lateral transfer in prokaryote genome evolution. Proc. Natl. Acad. Sci., 105, 10039–10044.

Dehoux, P. et al. (2016) Comparative genomics of Clostridium bolteae and Clostridium clostridioforme reveals species-specific genomic properties and numerous putative antibiotic resistance determinants. BMC Genomics, 17, 819.

Fodor, A.A. et al. (2012) The ‘most wanted’ taxa from the human microbiome for whole genome sequencing. PLoS One, 7.

Greenblum, S. et al. (2015) Extensive Strain-Level Copy-Number Variation across Human Gut Microbiome Species. Cell, 160, 583–594.

Holt, K.E. et al. (2015) Genomic analysis of diversity, population structure, virulence, and antimicrobial resistance in *Klebsiella pneumoniae*, an urgent threat to public health. Proc. Natl Acad. Sci., 112, E3574–E3581.

Huson, L.W. (2007) Performance of Some Correlation Coefficients When Applied to Zero-Clustered Data. J. Mod Appl. Stat. Methods, 6, 530–536.

Jie, Z. et al. (2017) The gut microbiome in atherosclerotic cardiovascular disease. Nat. Commun., 8, 845.

Jovel, J. et al. (2016) Characterization of the Gut Microbiome Using 16S or Shotgun Metagenomics. Front. Microbiol., 7.

Karlsson, F.H. et al. (2014) Metagenomic Data Utilization and Analysis (MEDUSA) and Construction of a Global Gut Microbial Gene Catalogue. PLoS Comput. Biol., 10.

Koonin, E. V. and Wolf, Y.I. (2008) Genomics of bacteria and archaea: The emerging dynamic view of the prokaryotic world. Nucleic Acids Res., 36, 6688–6719.

Kowalski, C.J. (1972) On the Effects of Non-Normality on the Distribution of the Sample Product-Moment Correlation Coefficient. Appl. Stat., 21, 1.

Kultima, J.R. et al. (2012) MOCAT: A Metagenomics Assembly and Gene Prediction Toolkit. PLoS One, 7.

Lagier, J.-C. et al. (2016) Culture of previously uncultured members of the human gut microbiota by culturomics. Nat. Microbiol., 1, 16203.

Larsbrink, J. et al. (2014) A discrete genetic locus confers xyloglucan metabolism in select human gut Bacteroidetes. Nature, 506, 498–502.

Li, J. et al. (2014) An integrated catalog of reference genes in the human gut microbiome. Nat. Biotechnol., 32, 834–841.

Lin, L.I.-K. (1989) A Concordance Correlation Coefficient to Evaluate Reproducibility. Biometrics, 45, 255.

Loman, N.J. et al. (2013) A Culture-Independent Sequence-Based Metagenomics Approach to the Investigation of an Outbreak of Shiga-Toxigenic Escherichia coli O104:H4. JAMA, 309, 1502.

Luo, C. et al. (2015) ConStrains identifies microbial strains in metagenomic datasets. Nat. Biotechnol., 33, 1045–1052.

Medini, D. et al. (2005) The microbial pan-genome. Curr. Opin. Genet. Dev., 15, 589–594.

Nayfach, S. et al. (2016) An integrated metagenomics pipeline for strain profiling reveals novel patterns of bacterial transmission and biogeography. Genome Res., 26, 1612–1625.

Nielsen, H.B. et al. (2014) Identification and assembly of genomes and genetic elements in complex metagenomic samples without using reference genomes. Nat. Biotechnol., 32, 822–828.

Osborne, J.W. and Overbay, A. (2004) The power of outliers (and why researchers should always check for them). Pract. Assessment, Res. Eval., 9, 1–8.

Qin, J. et al. (2012) A metagenome-wide association study of gut microbiota in type 2 diabetes. Nature, 490, 55–60.

Ranjan, R. et al. (2016) Analysis of the microbiome: Advantages of whole genome shotgun versus 16S amplicon sequencing. Biochem. Biophys. Res. Commun., 469, 967–977.

Scaria, J. et al. (2010) Analysis of Ultra Low Genome Conservation in Clostridium difficile. PLoS One, 5, e15147.

Schloissnig, S. et al. (2012) Genomic variation landscape of the human gut microbiome. Nature, 493, 45–50.

Scholz, M. et al. (2016) Strain-level microbial epidemiology and population genomics from shotgun metagenomics. Nat. Methods, 13, 435–438.

Schwartzman, A. and Lin, X. (2011) The effect of correlation in false discovery rate estimation. Biometrika, 98, 199–214.

Sczyrba, A. et al. (2017) Critical Assessment of Metagenome Interpretation - A benchmark of metagenomics software. Nat. Methods, 14, 1063–1071.

Sunagawa, S. et al. (2013) Metagenomic species profiling using universal phylogenetic marker genes. Nat. Methods, 10, 1196–1199.

Touchon, M. et al. (2009) Organised genome dynamics in the Escherichia coli species results in highly diverse adaptive paths. PLoS Genet., 5, e1000344.

Truong, D.T. et al. (2017) Microbial strain-level population structure & genetic diversity from metagenomes. Genome Res., 27, 626–638.

Větrovský, T. and Baldrian, P. (2013) The Variability of the 16S rRNA Gene in Bacterial Genomes and Its Consequences for Bacterial Community Analyses. PLoS One, 8.

Wang, J. and Jia, H. (2016) Metagenome-wide association studies: fine-mining the microbiome. Nat. Rev. Microbiol., 14, 508–522.

Wang, W.L. et al. (2015) Application of metagenomics in the human gut microbiome. World J. Gastroenterol., 21, 803–814.

Zhu, A. et al. (2015) Inter-individual differences in the gene content of human gut bacterial species. Genome Biol., 16, 82.

